# Tissue-specific safety mechanism results in opposite protein aggregation patterns during aging

**DOI:** 10.1101/2020.12.04.409771

**Authors:** Raimund Jung, Marie C. Lechler, Christian Rödelsperger, Waltraud Röseler, Ralf J. Sommer, Della C. David

## Abstract

During aging, proteostasis capacity declines and aggregation-prone proteins become instable, accumulating in protein aggregates both inside and outside cells. Both in disease and during aging, proteins selectively aggregate in certain tissues and not others. Yet, tissue-specific regulation of protein aggregation remains poorly understood. Surprisingly, we found that the inhibition of three core protein quality control systems, i.e. chaperones, proteasome and macroautophagy, leads to lower levels of age-dependent protein aggregation in *C. elegans* pharyngeal muscles, but higher levels in body-wall muscles. We describe a novel safety mechanism called SAPA that selectively targets newly synthesized aggregation-prone proteins to suppress aggregation and proteotoxicity. *vha-12* and *scav-3* mutants reveal that SAPA relies on macroautophagy-independent lysosomal degradation. Furthermore, SAPA involves several previously uncharacterized components of the intracellular pathogen response. We propose that SAPA represents an anti-aggregation machinery targeting aggregation-prone proteins for lysosomal degradation.

## Introduction

Active mechanisms to ensure functional and undamaged proteins are crucial across lifeforms [1]. A healthy proteome is maintained both inside and outside cells by a complex network of proteostasis components [2, 3]. However, with age, disruption of protein folding and proteolytic activities leads to a decline in proteostasis capacity [3]. Inhibition of protein quality control (PQC) systems and acute stress such as heat shock cause protein instability and ultimately protein aggregation [4–7]. Aberrant protein deposition with highly structured amyloid fibrils is a prominent pathological feature in agerelated diseases such neurodegenerative disorders, amyloidoses and type II diabetes [8]. Despite the potentially toxic nature of protein aggregation, certain physiological processes rely on discrete amyloid formation [9]. With the concomitant decline in proteostasis during aging, the intrinsic aggregation propensity of part of the proteome becomes a challenge for the organism [10–12] and widespread protein aggregation is found in aged animals of different species [5, 13–21]. Age-dependent aggregates show similarities to disease-associated protein aggregates and contain amyloid-like structures [18, 22]. Notably, evidence from the model organism *Caenorhabditis elegans* reveals that age-dependent aggregates are toxic and accelerate the functional decline of tissues [22, 23]. Moreover, age-dependent aggregates can initiate disease-associated protein aggregation [16].

Certain tissues and cell types are more likely to accumulate aggregates than others. In the early stages of neurodegenerative disorders, selective vulnerability to protein aggregation in distinct brain regions is particularly striking despite similar levels of expression of the disease-associated protein in less-susceptible brain regions [24, 25]. Emerging evidence highlights cell- and tissue-specificity also in the age-dependent aggregation process [15, 22]. However, the causes driving selective vulnerability or resilience to protein aggregation remain poorly understood [24, 25]. One possible explanation is that different cells and tissues employ divergent proteostasis strategies. In particular, evidence from *C. elegans* reveals that tissues rely to varying degrees on the chaperone system, proteasome or macroautophagy degradation to alleviate protein misfolding in an age-dependent manner [26–29]. In addition to conventional PQC, specialized mechanisms monitor the aggregation process, such as active sequestration of aggregation-prone proteins into compartments [30–32], disassembly by specialized chaperones [33–35], selective degradation of aggregates by macroautophagy (aggrephagy) [36] or expulsion in exophers [37].

Most of our current knowledge about the regulation of protein aggregation has been gained by examining disease-associated aggregating proteins as well as a few ectopically expressed standard aggregation-prone proteins such as luciferase. Expanding research into age-dependent protein aggregation and its regulation in different tissues offers the opportunity to identify novel mechanisms to counteract selective vulnerability to protein aggregation. Here, we report the discovery of a tissue-specific mechanism that actively prevents protein aggregation in *C. elegans* pharyngeal muscles but not in the body-wall muscles. The safety mechanism is triggered in response to defective conventional PQC and preferentially targets the aggregation process of globular proteins. We found that aggregation is effectively curbed by removing newly synthesized aggregation-prone proteins rather than clearing large aggregates. To prevent aggregation specifically in the pharynx, the safety mechanism relies on macroautophagy-independent lysosomal degradation and uncharacterized factors previously connected to the host’s response to natural intracellular pathogens affecting the digestive tract. Notably by limiting protein aggregation, the safety mechanism alleviates proteotoxicity.

## Results

### PQC impairment induces a protective safety mechanism in pharyngeal muscles

To identify potential tissue-specific mechanisms controlling age-dependent protein aggregation, we investigated the role of PQC in two different tissues in *C. elegans*: the pharyngeal non-striated muscles and the striated body-wall muscles. To evaluate changes in protein aggregation in response to PQC impairment, we examined fluorescent-tagged Ras-like GTP-binding protein rhoA (RHO-1) in pharyngeal muscles and Casein kinase I isoform alpha (KIN-19) aggregation in the pharynx and body-wall muscles. Both proteins are highly prone to aggregate with age and redistribute into compact puncta containing amyloid-like structures similar to disease-associated protein aggregates [18, 22]. We quantified changes in aggregation in response to knocking down the three canonical core PQC systems: chaperone-mediated folding, degradation by the proteasome and macroautophagy. Specifically, we targeted heat shock factor 1 (HSF-1), the main transcription factor controlling chaperone expression in *C. elegans*, proteasome subunits of the 20S core and 19S cap (PBS-3, PAS-6, RPT-6) and two essential macroautophagy components involved in the formation of autophagosome precursors (ATG-18, ortholog of human WIPI1 and WIPI2 as well as UNC-51, ortholog of human ULK1 and ULK2). To our surprise, we observed opposite outcomes for protein aggregation in the different muscle types: We observed that PQC inhibition accelerated KIN-19 aggregation with age in the body-wall muscles (Fig. 1 A and B). In contrast, PQC inhibition led to a significant reduction in RHO-1 and KIN-19 aggregation in the pharyngeal muscles in both young and aged animals (Fig. 1 C to F, Fig. S1 A to D and Data File S1). These results indicate that a tissue-specific safety mechanism in pharyngeal muscle cells is triggered in response to PQC inhibition to prevent protein aggregation, which we name safeguard against protein aggregation (SAPA).

**Figure 1:**
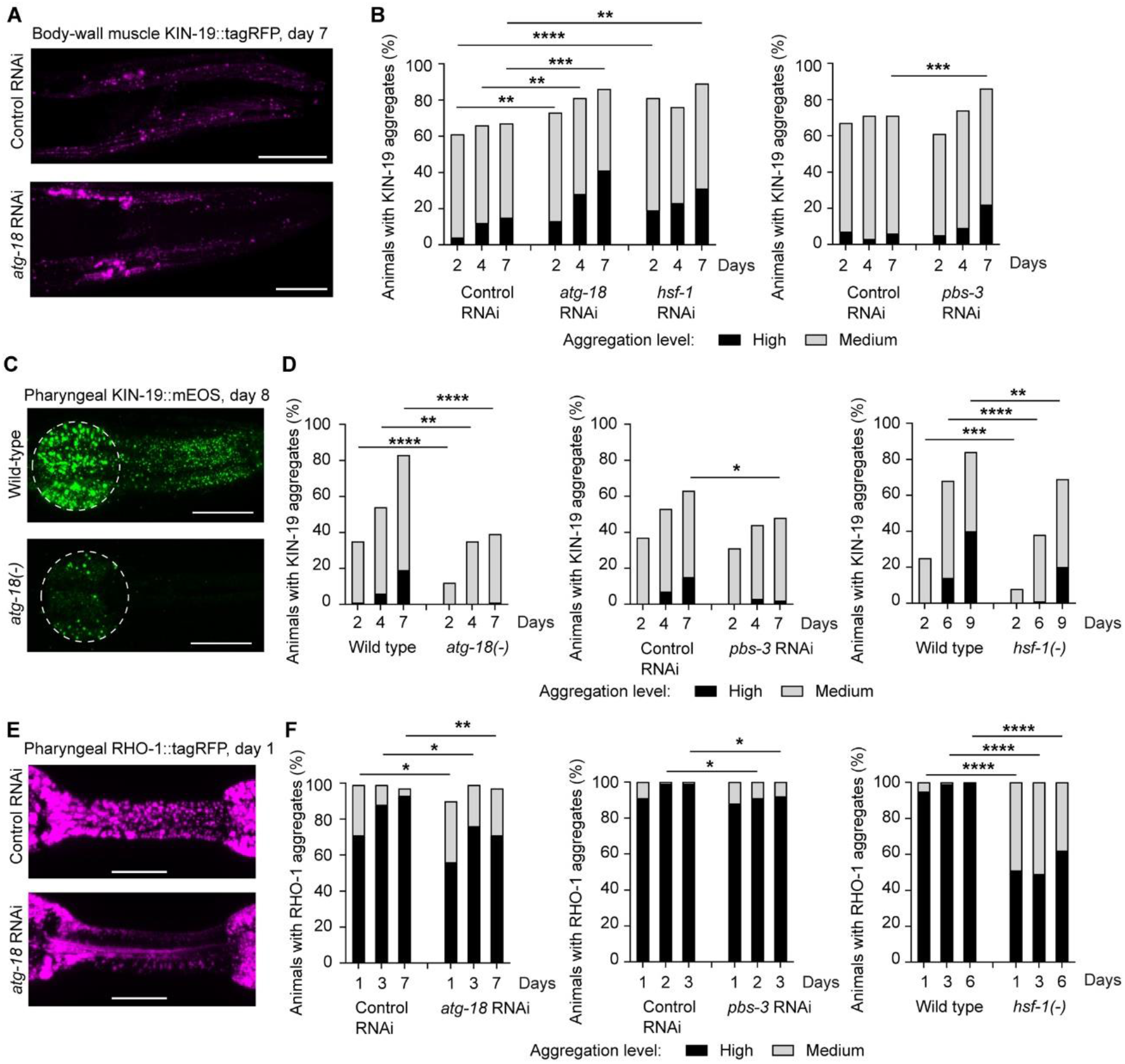
PQC impairment leads to reduced protein aggregation in pharyngeal muscles. (A, B) KIN-19 aggregation is increased in body-wall muscles upon PQC inhibition. Representative confocal images of KIN-19::tagRFP in the head body-wall muscles shown as maximum z-stack projections. Scale bar: 30 μm (A). Changes in KIN-19::tagRFP aggregation evaluated over time in the population with impaired macroautophagy (*atg-18*), impaired proteasomal degradation (*pbs-3*) and reduced chaperone levels (*hsf-1*) (B). (C, D) Aged animals with impaired PQC have less KIN-19::mEOS2 aggregates in the pharyngeal muscles. Representative confocal images of animals expressing KIN-19::mEOS2 in the pharynx shown as maximum z-stack projections with anterior pharyngeal bulb circled in white. Scale bar: 20 μm (C). Changes in KIN-19::mEOS2 aggregation evaluated over time in the population (D). (E, F) Young animals with impaired PQC have less RHO-1::tagRFP aggregates in the pharyngeal muscles. Representative confocal images of RHO-1::tagRFP in the pharyngeal isthmus shown as maximum z-stack projections. Scale bar: 15 μm (E). Changes in RHO-1::tagRFP aggregation evaluated over time in the population with impaired PQC (F). P-values determined by Fisher’s exact test, Chi-test and ordinal logistic regression. *p<0.05, **p<0.01, ***p<0.001, ****p<0.0001 See also Data File S1 for number of animals evaluated and statistics and Fig. S1.

### SAPA displays aggregation-prone substrate specificity

Next, we investigated whether SAPA is effective in preventing the aggregation of RNA-binding protein polyadenylate-binding protein 1 (PAB-1). PAB-1 contains a low complexity prion-like domain and undergoes liquid-liquid phase separation to form stress granules, putting it at risk of aggregating during aging or chronic stress in *C. elegans [23, 38]*. In contrast, both KIN-19 and RHO-1 are highly structured globular proteins, which are intrinsically prone to aggregate shortly after synthesis [22]. Notably, the divergent aggregation processes are controlled by different mechanisms [23, 38, 39]. Unlike KIN-19 or RHO-1 aggregation, PAB-1 aggregation in the pharynx was accelerated by inhibition of chaperones (as previously described [23]), proteasome and autophagy systems (Fig. S1E). These findings imply that SAPA displays substrate specificity towards aggregation-prone globular proteins.

### SAPA prevents de novo formation of aggregates

To gain insight into how SAPA regulates protein aggregation, we investigated the dynamics of aggregate formation and removal upon PQC failure. For this, we used KIN-19 and RHO-1 tagged with mEOS2, a green-to-red photoconvertible fluorescent protein [40]. By exposing the animals to intense blue light, all aggregates present were converted to emitting red fluorescence. Of note, the core region of some large aggregates was refractory to photoconversion (Fig. 2 and as previously described [22]). Thereafter, we followed the rate of new (green emitting) aggregate formation and the rate of old (red emitting) aggregate removal over time. During aging, KIN-19 and RHO-1 are prone to form aggregates shortly after synthesis while old aggregates are slowly removed (Fig. 2 A to C) [22]. Upon PQC impairment, the rate of old aggregate removal was either similar for KIN-19 aggregates or moderately enhanced for RHO-1 aggregates compared to control conditions (Fig. 2 A to C). In contrast, we observed a striking effect on the formation of new aggregates. During proteasome or macroautophagy disruption, the percentage of animals with new KIN-19 and RHO-1 aggregates gained over 48 hours was strongly reduced compared to control conditions (Fig. 2 A to C). In the body-wall muscles where we did not observe activation of a safety mechanism (Fig. 1 A and B), we found no difference between the rate of new aggregate formation and old aggregate removal when impairing proteasome activity compared to control conditions (Fig. S2A). Together, these results reveal that SAPA mainly limits the formation of new aggregates in the pharynx.

**Figure 2:**
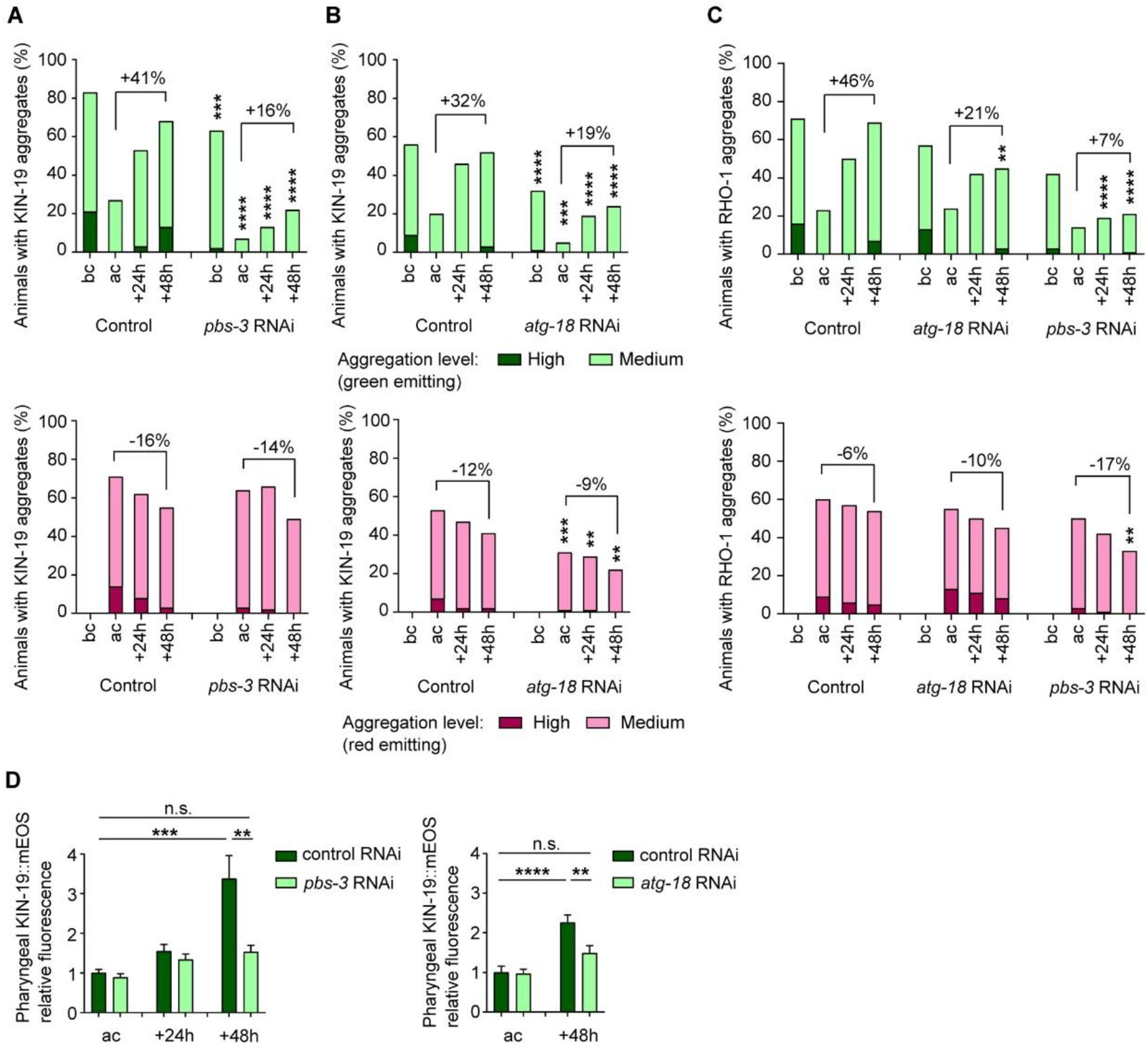
SAPA prevents de novo formation of aggregates in the pharynx by promoting removal of newly synthesized aggregation-prone protein. (A, B) Formation of new green-labeled KIN-19::mEOS2 aggregates is delayed in the pharynx of animals with impaired protein degradation whereas the removal of old red-labeled aggregates is not affected. Proteasome inhibition by *pbs-3* RNAi (A) and macroautophagy inhibition by *atg-18* RNAi (B). (C) Formation of new green-labeled RHO-1::mEOS2 aggregates is delayed in the pharynx of animals with impaired protein degradation. (A to C) Conversion performed at day 5. Before conversion (bc), after conversion (ac). In graph, difference in percent of animals with aggregation between 48h and after conversion. (D) After photoconversion of pharyngeal KIN-19::mEOS2 in young day 2 adults, monitoring of total green fluorescence reveals reduced accumulation over time of newly synthesized aggregation-prone protein upon proteasome impairment or macroautophagy impairment. Relative fluorescence represents total fluorescence of treatment normalized to control RNAi after conversion. SEM shown. P-values determined by Fisher’s exact test comparing treatment versus control at the respective time point (A to C) and unpaired T-test (D). **p<0.01, ***p<0.001, ****p<0.0001 See also Data File S1 for number of animals evaluated and statistics and Fig. S2.

### SAPA promotes removal of newly synthesized aggregation-prone proteins

To understand how SAPA interferes with the formation of new aggregates, we quantified changes in total levels of green-emitting KIN-19 upon PQC impairment in young animals during the early stages of protein aggregation. If SAPA hinders the final assembly step into large visible aggregates, we would expect an accumulation of diffuse newly synthesized aggregation-prone proteins over time. In contrast, if SAPA prevents aggregation by removing newly synthesized aggregation-prone proteins, we would expect less green-emitting KIN-19 to accumulate over time. In support of the latter, SAPA stopped the accumulation of newly synthesized KIN-19 over 48 hours in response to proteasome and macroautophagy inhibition (Fig. 2D). In control animals expressing the fluorescent protein mEOS2 alone, which does not aggregate [22], there was no reduction in newly synthesized protein in response to PQC impairment (Fig. S2B). These control results support the specificity of SAPA and exclude a general decrease in translation. Of note, overall higher levels of mEOS2 after 48 hours is likely due to enhanced *kin-19* promoter activity with age as previously described [18]. Together, these results imply that SAPA targets newly synthesized aggregation-prone proteins for removal in order to avoid de novo formation of aggregates upon PQC disruption.

### Macroautophagy-independent lysosomal degradation protects against protein aggregation

Next, we examined how SAPA promotes the removal of aggregation-prone proteins. SAPA could be due to a compensatory upregulation of one of the degradation systems [41, 42]. However, combining proteasome and macroautophagy inhibition or HSF-1 and macroautophagy inhibition failed to restore protein aggregation (Fig. 3 A and B). In addition to macroautophagy, other forms of autophagy can target cytoplasmic material for lysosomal degradation, such as general microautophagy and chaperone-mediated autophagy [43, 44]. Yet, it is unclear whether these types of autophagy occur in *C. elegans*. To investigate the role of the lysosome, we used loss-of-function mutants for the vacuolar proton translocating ATPase (VHA-12 subunit) responsible for lysosome acidification and the lysosomal membrane protein SCAV-3, a regulator of lysosome integrity [45]. Of note, SCAV-3 is prominently expressed in the pharynx. We found that impairing lysosomal degradation through either *vha-12* or *scav-3* loss-of-function eliminated SAPA triggered by macroautophagy inhibition and restored RHO-1 aggregation to levels seen in control conditions without PQC disruption (Fig. 3 C and D). Thus, SAPA depends on macroautophagy-independent lysosomal degradation to limit protein aggregation.

**Figure 3:**
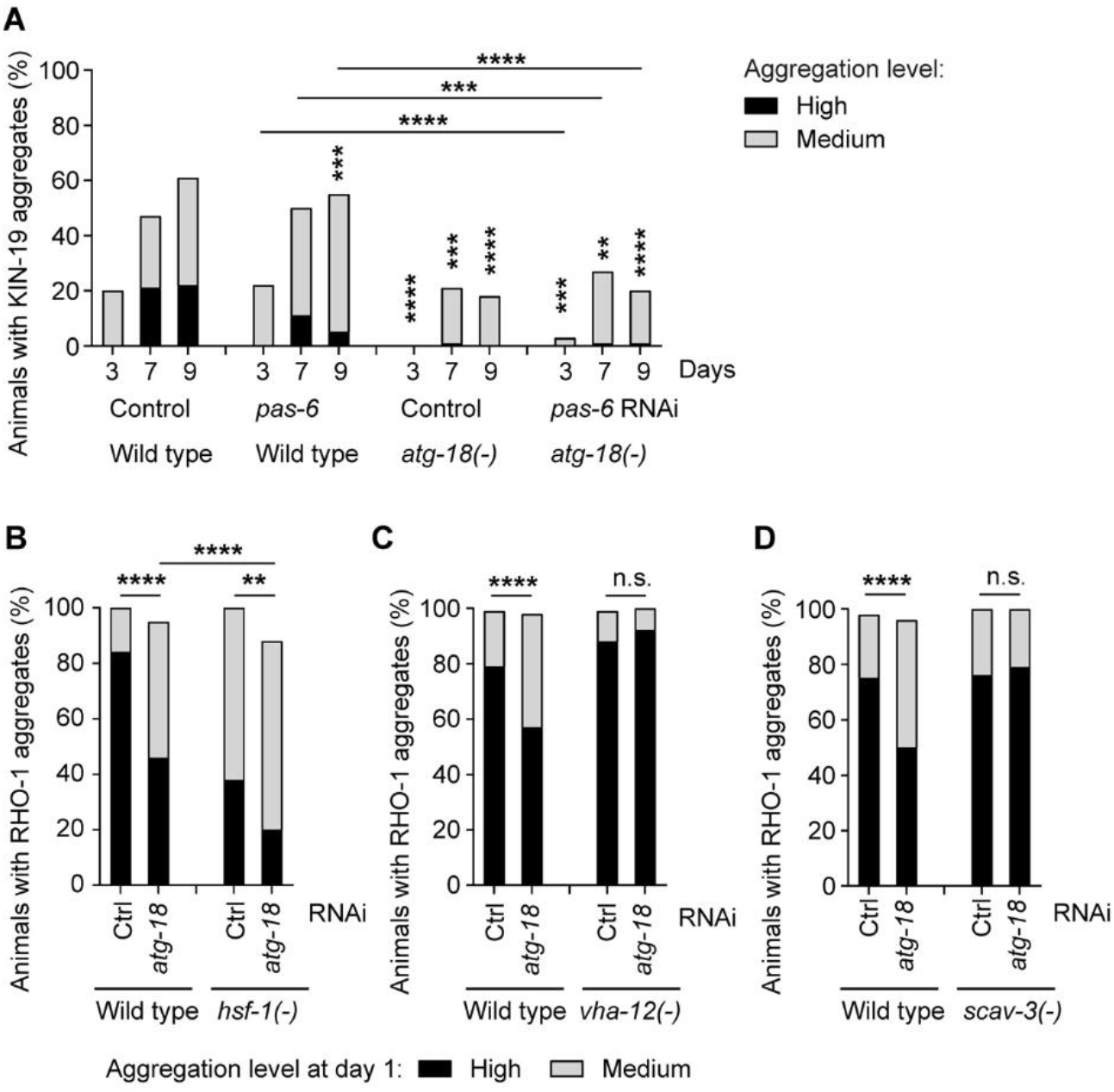
Macroautophagy-independent lysosomal degradation is responsible for preventing protein aggregation. (A) Reduced pharyngeal KIN-19 aggregation upon proteasome inhibition (*pas-6* RNAi) is not restored by macroautophagy inhibition (*atg-18(-)*). Changes in KIN-19::mEOS2 aggregation evaluated over time in the worm population. (B) Reduced pharyngeal RHO-1 aggregation upon chaperone impairment (*hsf-1(-)*) is not restored by macroautophagy inhibition (*atg-18* RNAi). Changes in RHO-1::tagRFP aggregation at day 1 in the worm population. (C, D) Reduced pharyngeal RHO-1 aggregation upon macroautophagy impairment (*atg-18* RNAi) is restored by impairing lysosomal degradation (*vha-12(-)* (C) and (*scav-3(-)* (D)). Changes in RHO-1::tagRFP aggregation at day 1 in the worm population. P-values determined by Fisher’s exact test and Chi-test. **p<0.01, ***p<0.001, ****p<0.0001 See also Data File S1 for number of animals evaluated and statistics.

### SAPA relies on components of the intracellular pathogen response

The question remains: why is SAPA tissue-specific? To address this, we performed RNA sequencing to identify expression changes selectively induced by PQC impairment in animals with pharyngeal protein aggregates but not induced by PQC impairment in animals with body-wall muscle aggregates (Fig. S3, Data File S2). With this experimental design, we identified 12 genes significantly upregulated that could play a role in SAPA (Table S1, Data File S2). Among these, two genes have human orthologs: C01B10.4 has sequence similarities with Butyrylcholinesterase (E-value: 4e-30) and C53A5.11 has sequence similarities with Actin-binding protein IPP (E-value: 1.2e-27). Notably, five out of the 12 candidates for SAPA are genes of unknown function previously identified as part of the host’s intracellular pathogen response (IPR) (Data File S2) [46]. We targeted eight of these 12 genes by RNAi. Upon PQC failure, knockdown of six out of eight genes reproducibly contributed to restoring RHO-1 aggregation in the pharynx of *hsf-1* mutants (Fig. 4A, Table S1 and Data File S3), confirming their participation in SAPA. In contrast in the wild-type background, the RNAi treatments had no effect on RHO-1 aggregation (Fig. 4B and Data File S3), highlighting that PQC impairment is required to trigger SAPA. Among the SAPA components discovered by RNA sequencing, PALS-5 is one of the most highly upregulated genes in response to intracellular pathogens [46]. Overexpressed PALS-5 consistently co-localized with RHO-1 aggregates, suggesting an interaction between both proteins (Fig. 4C). In contrast, we observed co-localization of PALS-5 with only a few PAB-1 aggregates (Fig. 4D). Using a GFP transcriptional reporter for *pals-5*, we found expression of PALS-5 in the pharynx and intestine but not in the body-wall muscles (Fig. S4 A to D). Importantly, overexpression of PALS-5 in the pharynx prevented RHO-1 aggregation but not PAB-1 aggregation (Fig. 4 E to G). Thus, PALS-5 on its own is sufficient to limit protein aggregation, suggesting that the local upregulation of a select group of IPR factors is an essential part of SAPA.

**Figure 4:**
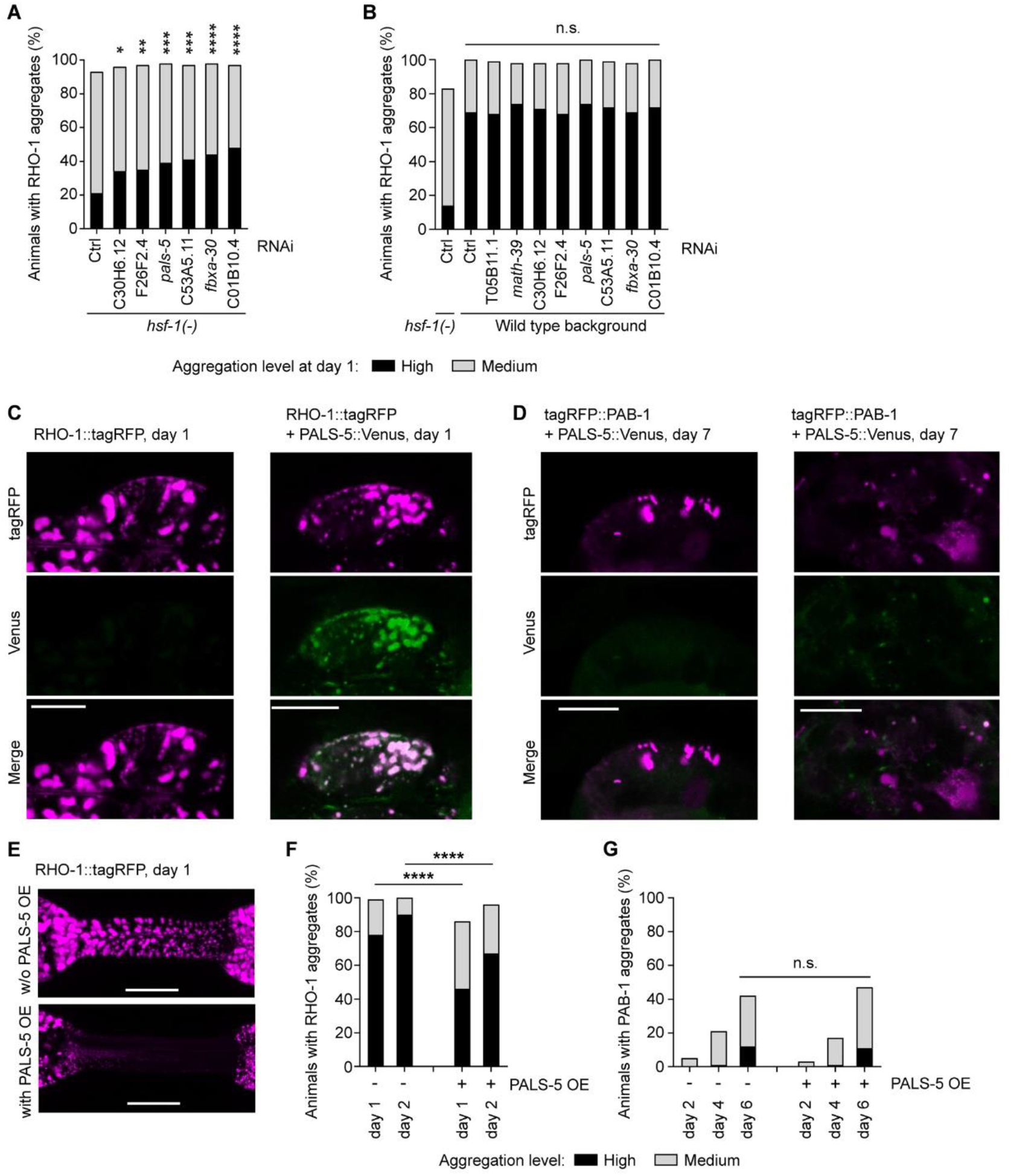
SAPA relies on factors involved in the intracellular pathogen response. (A, B) Knockdown by RNAi of genes identified by RNA-seq as selectively upregulated by PQC failure in transgenic animals with pharyngeal RHO-1 and KIN-19 aggregation. Quantification of RHO-1::tagRFP aggregation at day 1 with PQC failure induced by *hsf-(−)* (A) and without PQC failure (B). (C, D) Representative confocal single plan images of pharyngeal RHO-1::tagRFP (magenta, 0.5% laser), PALS-5::Venus (green, 5% laser), tagRFP::PAB-1 (magenta, 10% laser), scale bar 10 μm. PALS-5 colocalizes with RHO-1 aggregates (right panel) compared to no signal in the absence of PALS-5 OE (left panel) (C). No co-localization (left panel) or partial co-localization of PALS-5 with a few PAB-1 aggregates (right panel) (D). (E, F) PALS-5 overexpression delays RHO-1 aggregation in the pharynx. Representative confocal images of RHO-1::tagRFP in the pharyngeal isthmus in animals without (w/o) PALS-5 overexpression (OE) (top) and with PALS-5 OE (below), shown as maximum z-stack projections. Scale bar: 15 μm (E). Quantification of RHO-1::tagRFP aggregation in the worm population (F). (G) PALS-5 overexpression does not influence PAB-1 aggregation in the worm population. Ordinal logistic regression analysis comparing all RNAi treatments to the control RNAi (A, B), Fisher’s exact test and Chi-square test comparing same days with and without PALS-5 OE (F, G). *p<0.05, **p<0.01, ***p<0.001, ****p<0.0001 See also Data File S1 for number of animals evaluated and statistics, Data File S2 for RNA-seq data, Data File S3 for repeats, Fig. S3 and S4.

### SAPA lessens proteotoxicity

Finally, we asked whether preventing protein aggregation during PQC failure confers a physiological advantage to the organism. It was previously shown that age-dependent protein aggregates accelerate age-related functional decline [22]. Moreover, RHO-1 aggregates present in the pharyngeal muscles of young animals are sufficient to impair pharyngeal pumping, demonstrating that these aggregates are toxic independent of aging processes. Here, we evaluated how lowering RHO-1 aggregation through SAPA affects the pumping phenotype. To induce SAPA, we used HSF-1 inhibition with the hypomorphic *hsf-1* mutation, which leads to a strong reduction in RHO-1 aggregation in young animals (Fig. 1F). Compared to inhibition of macroautophagy or proteasome activity, these mutants are relatively healthy [47]. Importantly, we found that *hsf-1* inhibition partially averted the decline in pharyngeal pumping caused by RHO-1 aggregation (Fig. 5 A to C). Thus, effectively preventing the process of aggregation during PQC failure is beneficial to the organism at the physiological level.

**Figure 5:**
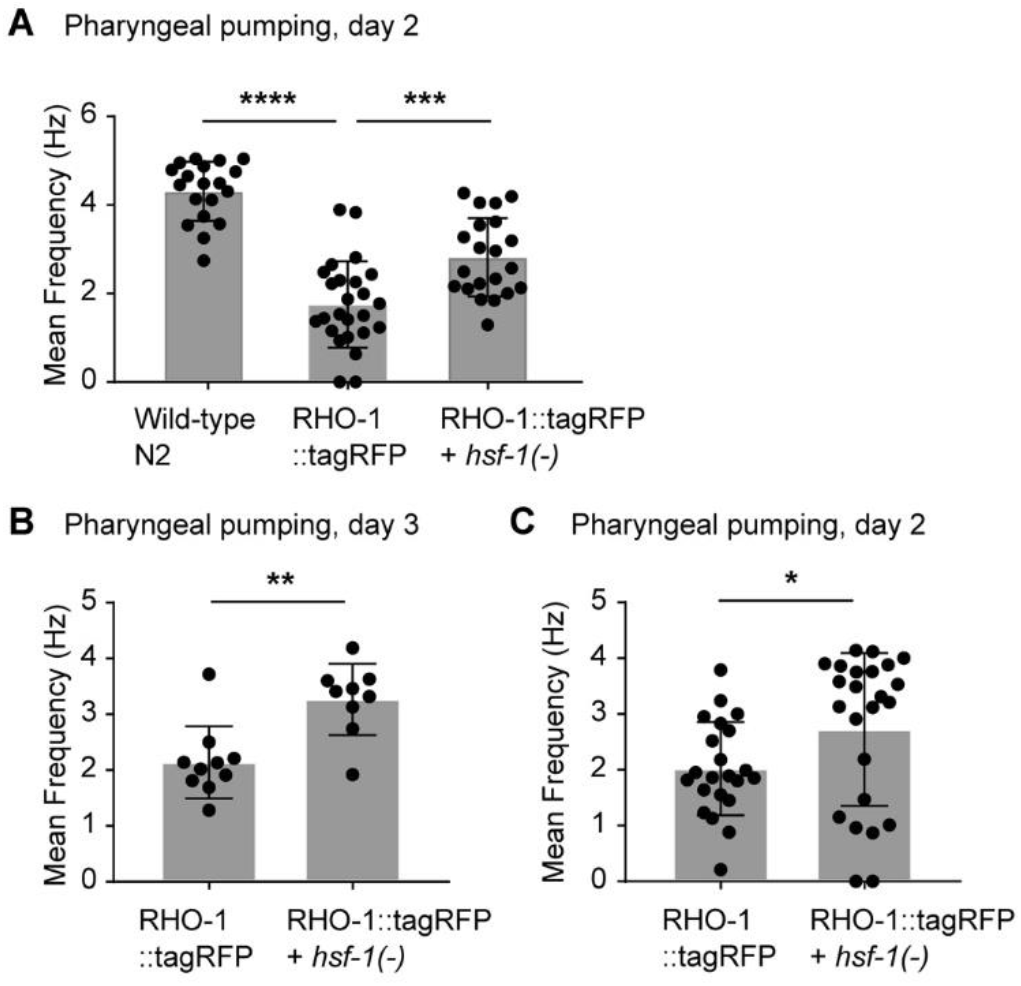
Reducing RHO-1 aggregation alleviates proteotoxicity. (A, B, C) *hsf-1* mutants rescue pharyngeal pumping defects caused by RHO-1 aggregation. Each dot represents mean frequency measured in individual animals. Histogram represents mean of all measurements per condition with SD. One-way ANOVA with Tukey’s multiple comparisons test (A) and Unpaired T-test (B, C). *p<0.05, **p<0.01, ***p<0.001, ****p<0.0001 See also Data File S1 for number of animals evaluated and statistics.

## Discussion

Aberrant uncontrolled protein aggregation is toxic to the organism as observed in neurodegenerative diseases and amyloidosis as well as during aging [8, 22]. Enhancing proteostasis is a promising strategy to counteract proteotoxicity and considered in efforts to develop an effective therapy for diseases of protein aggregation [48]. In this current study, we used intrinsically aggregation-prone proteins in *C. elegans* to understand the proteostasis of age-dependent protein aggregation. We describe a novel tissue-specific safety mechanism triggered in response to PQC impairment that limits protein aggregation and its toxicity.

Our findings reveal that the safety mechanism does not target all aggregation-prone substrates equally. SAPA was highly effective in preventing the aggregation of two normally globular proteins, casein kinase I isoform alpha (KIN-19) and ras-like GTP-binding protein rhoA (RHO-1). In contrast, SAPA had no effect on the aggregation of a stress granule marker, polyadenylate-binding protein 1 (PAB-1). As the aggregation of both RHO-1 and KIN-19 starts shortly after synthesis [22] and as SAPA limits new aggregate formation (Fig. 2), it is likely that either unfolded states or early intermediate aggregation species are specifically recognized for SAPA mediated degradation. Unlike KIN-19 and RHO-1, PAB-1 contains a low-complexity prion-like domain and undergoes liquid-liquid phase separation to form stress granules [23]. In vitro and in vivo work demonstrate that solidification into aggregates can be initiated in the context of the liquid droplet [38, 49]. Thus, the selectivity of SAPA may be explained by differences in the aggregation process of stress granule proteins compared to non-stress granule components. This is in agreement with previous work showing different strategies used by long-lived *C. elegans* with reduced insulin/IGF-1 signaling to promote preferentially the solubility of stress granule proteins compared to other types of aggregation-prone proteins [23].

We found that SAPA is triggered in response to the perturbation of key systems maintaining protein homeostasis, namely protein folding by chaperones and protein degradation by the proteasome and macroautophagy. SAPA is likely distinct from previously identified compensatory mechanisms activated in response to PQC impairment. Particularly well characterized is the upregulation of macroautophagy in response to proteasome defects to ensure the removal of excessive levels of ubiquitinated proteins [41, 42]. Yet, we found that SAPA is still functional when inhibiting both macroautophagy and proteasome-mediated degradation (Fig. 4 A). In response to macroautophagy impairment, golgi membrane-associated degradation (GOMED) is induced to compensate [50]. SAPA is likely also distinct from GOMED as SAPA is triggered by *unc-51* inhibition whereas GOMED is dependent on Ulk-1 (homolog of *C. elegans unc-51*). Removal of protein aggregates by expulsion from neurons through exopher formation is enhanced in *C. elegans* during proteasome, macroautophagy or *hsf-1* inhibition [37]. Yet it is unlikely that exopher formation greatly contributes to protecting pharyngeal muscle cells from age-dependent protein aggregation, the pharynx being isolated from the pseudocoelom by a thick basal lamina. Instead, our data reveal that in response to core PQC failure, newly-synthesized aggregation-prone proteins are cleared before forming large aggregates by non-canonical degradation. While macroautophagy is the best characterized form of autophagy, cytosolic proteins can be targeted to the lysosome through other forms of autophagy. Chaperone-mediated autophagy specifically targets proteins for import into the lysosome through HSC70 and LAMP2A [51]. However, no ortholog of LAMP2A has been identified in *C. elegans* [52]. Another possibility to degrade cytosolic proteins without using autophagosomes is general microautophagy where the cargo is taken up by vesicles formed at the surface of the lysosome or by late endosomes [43, 44]. Microautophagy has yet to be characterized in *C. elegans* [52], yet, two recent studies provide evidence for macroautophagy-independent degradation of cytosolic cargo by the lysosome evocative of microautophagy *[53, 54]*. Further work is needed to establish how SAPA may rely on microautophagy to stop the aggregation of globular proteins during core PQC failure.

Selective vulnerability to protein aggregation remains a conundrum. Underlying genetic risk factors involved in protein degradation have been associated with major neurodegeneration disorders such as Alzheimer’s disease, Parkinson’s disease and Amyotrophic lateral sclerosis [25]. Moreover, increasing evidence highlights the importance of lysosomes in diseases of protein aggregation [55]. Yet, differences in aggregation-prone protein turnover between cell types [56] is probably determined by tissue-specific secondary proteostasis regulators [57] rather than differences in the core components of the degradative systems, which are ubiquitously expressed. In *C. elegans*, our analysis shows that the tissue specificity of SAPA is likely related to the expression of genes enriched in uncharacterized IPR factors. Overexpression of PALS-5, a commonly used reporter for IPR with unknown function, was sufficient to limit protein aggregation. Moreover human butyrylcholinesterase, the potential ortholog of SAPA component C01B10.4, inhibits amyloid-β aggregation *in vitro* [58]. Protein-protein interaction domains such as F-box and MATH (meprin-associated Traf homology) are overrepresented in IPR genes [46, 59] and we found three F-box domains and one MATH domain coding genes among the SAPA components. Interestingly, a number of F-box proteins act as adapters in ubiquitin-mediated protein degradation [60]. Thus, an intriguing possibility is that the SAPA components identified are anti-aggregation factors selectively targeting aggregation-prone proteins for degradation by lysosomes localized in the pharynx.

The clearance rate of disease-associated aggregating proteins is correlated with neuronal survival [56]. Similarly, we found that the safety mechanism triggered by PQC failure alleviated proteotoxicity and partially restored the pharyngeal pumping activity in *C. elegans*. Reduced proteotoxicity supports our findings that SAPA prevents aggregate formation by removing aggregation-prone proteins rather than allowing toxic intermediate aggregation species to accumulate. Why does the organism choose to prioritize resources to protect foremost the pharynx rather than body-wall muscles during PQC failure? The reason could be the fundamental role of the pharynx in feeding and its direct exposure to natural pathogens. The pharynx is a contractile organ that pumps bacterial food from the mouth of the worm into its intestine. By grinding up bacteria, the pharynx limits bacterial colonialization and infection in the intestine. Importantly, bacterial infection in the pharynx is associated with early death [61]. The digestive tract is also the point of entry for obligate intracellular pathogens such as viruses. Upon infection, the host triggers the IPR transcriptional program, which is distinct from the response to extracellular pathogens [46, 62, 63]. One of the host’s defense strategies against intracellular and extracellular pathogens is to improve resilience by enhancing proteostasis [2, 46, 62, 64]. Our work reveals that upon failure of PQC the organism co-opts part of the response to intracellular pathogens by inducing IPR genes such as *pals-5* to protect the pharynx against excessive proteostress. Consistent with this, *pals-5* expression in the digestive tract is induced by heat stress and proteasome inhibition as well as by intracellular pathogens [46, 62]. Whether these select IPR genes used by SAPA protect also against endogenous protein aggregation during an infection remains to be determined.

In conclusion, our findings characterize a novel tissue-specific safety mechanism that suppresses age-dependent protein aggregation and its toxicity in response to failure of core PQC systems. A better understanding of selective targeting of aggregation-prone proteins to the lysosome should help to design strategies to restore proteome health in vulnerable aged tissues.

## Materials and Methods

### Strains

Wild type: N2

Transgenics:

CF3166: *muEx473[pkin-19::kin-19::tagrfp + Ptph-1::gfp]*
CF3317: N2; *muEx512[Pkin-19::tagrfp + Ptph-1::gfp]*
CF3649: N2; *muIs209[Pmyo-3::kin-19::tagrfp + Ptph-1::gfp]*
CF3706: N2; *muEx587[Pkin-19::kin-19::meos2 + Punc-122::gfp]*
DCD13: N2; *uqIs9[Pmyo-2::rho-1::tagrfp + Ptph-1::gfp]*
DCD69: N2; *uqEx4[Pmyo-3::kin-19::meos2]*
DCD83: *ttTi5605II; unc-119(ed3)III; uqEx11[Pmyo-2::rho-1::meos2 + Punc-122::gfp + cb-unc-119(+)]*
DCD92: *hsf-1(sy441)I; uqIs9[Pmyo2::rho1::tagrfp + Ptph-1::gfp]*
DCD173: *hsf-1(sy441)I; muEx587[Pkin-19::kin-19::meos2 + Punc-122::gfp]*
DCD174: *atg-18(gk378)V; muEx587[Pkin-19::kin-19::meos2 + Punc-122::gfp]*
DCD214: N2; *uqIs24[Pmyo-2::tagrfp::pab1]*
DCD245: N2; *uqEx49[Pkin-19::meos2]*
DCD249: *hsf-1(sy441)I; uqIs24[Pmyo-2::tagrfp::pab1]*
DCD258: *hsf-1(sy441)I; muIs209[Pmyo-3::kin-19::tagrfp + Ptph-1::gfp]*
DCD324: *unc-51(e369)V; uqIs9[Pmyo2::rho1::tagrfp + Ptph-1::gfp]*
DCD326: *vha-12(ok821)X; uqIs9[Pmyo2::rho1::tagrfp + Ptph-1::gfp]*
DCD340: *unc-51(e369)V; uqIs24[Pmyo-2::tagrfp::pab1]*
DCD345: *scav-3(qx193)III; uqIs9[Pmyo2::rho1::tagrfp + Ptph-1::gfp]*
ERT54: *jyIs8 [Ppals-5::GFP + Pmyo-2::mcherry] X*
DCD411: N2; *uqIs9[Pmyo2::rho1::tagrfp + Ptph-1::gfp]; uqEx64[Pkin-19::pals-5::mVenus::histag + Punc-122::gfp]*
DCD415: N2; *uqIs24[Pmyo-2::tagrfp::pab1]; uqEx64[Pkin-19::pals-5::mVenus::histag + Punc-122::gfp]*

### Cloning and strain generation

Cloning and strain generation were performed as previously described [22]. Construct *Pkin-19::pals-5::mVenus::histag* and coinjection marker *Punc-122::gfp* were injected at 50 ng μl^−1^ each. Genetic crosses were made to transfer transgenes to the appropriate genetic background. The presence of the mutant allele was verified by polymerase chain reaction (PCR).

### *C. elegans* maintenance and RNAi knockdown

All strains were kept at 15°C on NGM plates inoculated with OP50 using standard techniques. Age-synchronization was achieved by transferring adults of the desired strain to 20°C and selecting their progeny at L4 stage. Adult mutants with *atg-18(-)* and *unc-51(-)* were kept at 15°C for laying eggs. From L4 stage, all experiments were performed at 20°C. RNAi treatment was performed by feeding as previously published [65]. The RNAi clones were acquired from the Marc Vidal or the Julie Ahringer RNAi feeding library (Source BioScience, UK) and sequenced. HT115 containing the empty vector L4440 was used as control. To start RNAi treatment from egg stage, adults were allowed to lay eggs on bacterial lawn expressing the dsRNA. To avoid developmental defects, proteasome subunits were targeted by RNAi from L4 stage. Day 1 of adulthood starts 24 hours after L4. To enhance the RNAi effects, F2 generation from animals subjected to RNAi treatment were evaluated when indicated.

### Imaging

For confocal analysis using a Leica SP8 confocal microscope with the HC PL APO CS2 63x/1.30 NA glycerol objective, worms were mounted onto slides with 2% agarose pads using 2 μM levamisole for anesthesia. For body-wall muscle images, worms were fixed in 4% paraformaldehyde (PFA) for 10 min at room temperature and imaged with the HC PL APO CS2 63x/1.40 NA oil objective. Leica HyD hybrid detector was used to detect KIN-19::mEOS2 (excitation: 506 nm, emission: 508 to 550 nm), PALS-5::Venus (excitation: 515 nm, emission: 521 to 545 nm), tagRFP::PAB-1 (excitation: 555 nm, emission: 565 to 650 nm). Leica PMT detector was used to detect RHO-1::tagRFP (excitation: 555 nm, emission: 565 to 620 nm).

### Photoconversion of mEOS2-tag and quantification of fluorescence levels

Photoconversion was performed as previously described by illuminating worms placed on a plate with blue fluorescence (387/11 BrightLine HC, diameter 40 mm) [22]. For all conversions, transgenic animals were exposed to blue light four times for five minutes, with 2 min pauses between exposures except for transgenics expressing *pmyo-2::RHO-1::mEOS2*, which were exposed to blue light five times for six minutes, with 2 minute pauses. For quantification of fluorescence levels, worms were mounted onto slides with 2% agarose pads using 2 μM levamisole for anesthesia. Using an Axio Observer Z1 (Zeiss), levels of green fluorescence (eGFP set 38HE, excitation 470 ±40 nm, emission 525 ±50 nm) were detected. Quantification of fluorescence levels was determined using ImageJ [66, 67] with the oval-shaped selection tool placed around the anterior bulb. Total fluorescence was obtained by subtracting the mean intensity in the anterior bulb from the mean background intensity and multiplying by the area measured.

### Aggregation quantification in vivo

Aggregation levels were determined using Leica fluorescence microscope M165 FC with a Planapo 2.0x objective. Aggregation was quantified following pre-set criteria adapted to the transgene expression pattern and levels in the different transgenic *C. elegans* models: Animals expressing *Pkin-19::KIN-19::mEOS2* or *Pkin-19::KIN-19::TagRFP* were divided into less than 10 puncta (low aggregation), between 10 and 100 puncta (medium aggregation) and over 100 puncta in the anterior pharyngeal bulb (high aggregation) [18, 22]. Because of extensive RHO-1 aggregation in young animals overexpressing *Pmyo-2::RHO-1::TagRFP* or *Pmyo-2::RHO-1::mEOS2*, aggregation was only quantified in the isthmus: animals with no aggregation (low aggregation), animals with aggregation in up to 50% (medium aggregation) and animals with aggregation in more than 50% (high aggregation) of the isthmus. Animals overexpressing *Pmyo-2:tagRFP:PAB-1* were divided into less than 10 puncta (low aggregation) and over 10 puncta (medium aggregation) in the posterior bulb and over 10 puncta in the anterior bulb (high aggregation) [23]. Animals overexpressing *Pmyo-3::KIN-19::TagRFP* were divided into over 15 puncta in the head or the middle body region (low aggregation), over 15 puncta in the head and the middle body region (medium aggregation) and over 15 puncta in head, middle body and tail region (high aggregation). The same categories defined for animals overexpressing *Pmyo-3::KIN-19::TagRFP* were used to evaluate animals overexpressing *Pmyo-3::KIN-19::mEOS2* with a cutoff of 10 puncta instead of 15 to account for slightly lower aggregation levels. Counting was done in a blind fashion for all conditions.

### Animal collection for RNA sequencing

Adults were allowed to lay eggs during 7 hours at 20°C on bacterial lawn expressing dsRNA for *atg-18*, *hsf-1* and empty vector L4440. To synchronize the animals, progeny at L4 stage were selected and kept at 20°C until collection. Animals overexpressing *Pmyo-2::RHO-1::TagRFP* (DCD13) were collected at day 2, 300 worms per condition. Animals overexpressing *Pkin-19::KIN-19::TagRFP* (CF3166) were collected at day 7, 230 worms per condition. Control animals overexpressing *Pmyo-3::KIN-19::TagRFP* (CF3649) were collected at day 4, 300 worms per condition. Control animals overexpressing *Pkin-19::TagRFP* (CF3317) were collected at day 7, 300 worms per condition. Collected animals were washed twice in M9 and snap frozen.

### RNA sequencing

Total RNA was isolated using Direct-zol RNA Mini-Prep Kit following the manufacturers’ instructions (Zymo Research, CA, USA). RNA concentration was quantified using Qubit (Invitrogen Life technologies, CA, USA) and Nanodrop (PEQLAB Biotechnologie GmbH, Erlangen, Germany) measurements. RNAseq libraries were prepared using TruSeq RNA library preparation kit v2 (Illumina Inc, CA, USA) according to the manufacturer’s instructions from 1 μg of total RNA in each sample. Libraries were quantified using Qubit and Bioanalyzer measurements (Agilent Technologies, CA, USA) and normalized to 2.5 nM. Samples were sequenced as 150bp paired end reads on multiplexed lanes of an Illumina HiSeq3000 (Illumina Inc, CA, USA). All sequencing data has been submitted to the European Nucleotide archive under the study accession PRJEB41493.

### Analysis of RNA-seq data

Raw RNA-seq reads were aligned to the *C. elegans* reference genome (Wormbase release WS250) with the help of tophat2 (version 2.0.14, default options) [68]. We used *C. elegans* gene annotations (Wormbase release WS250) to quantify expression levels and to perform tests for differential expression for all pairwise comparisons of samples with the software cuffdiff (version 2.2.1, default options) [69]. For further analysis, any gene with a P-value < 0.05 was considered as candidate for differential expression.

### Pharyngeal pumping analysis

Electrical activity of the pharyngeal pumping was measured using the NemaMetrix ScreenChip System (NemaMetrix, Eugene, OR) as previously described [22].

### Statistical analysis

For analysis of aggregation quantification in vivo, two-tailed Fisher’s exact test and Chi-test were performed with GraphPad. Fisher’s exact test was used for comparisons with two categories of aggregate levels where two of the three categories were combined together when a category counted less than five animals. For analysis of changes in fluorescence intensities, Student t-Test with two-tail distribution and two-sample unequal variance was performed with Excel. To analyze the effect on protein aggregation of multiple RNAi treatments compare to control RNAi, we used an ordinal logistic regression model, which was performed using R and its MASS package. Enrichment analysis of SAPA components in Data File 2 was performed with WormExp (http://wormexp.zoologie.uni-kiel.de/wormexp/) [70], category Microbes, one-sided Fisher’s exact test with Bonferroni correction P < 0.05.

## Acknowledgments

We thank X. Wang for sharing *scav-3(qx193)* mutants, C. Kenyon for providing some *C. elegans* strains and M. Schölling for help with statistics based on the ordinal logistic regression model. We thank E. Troemel for advice and critical input. Some strains were provided by the CGC, which is funded by NIH Office of Research Infrastructure Programs (P40 OD010440). This work was supported by funding from the DZNE, a Marie Curie International Reintegration Grant (322120 to DCD) and the Deutsche Forschungsgemeinschaft (DFG, German Research Foundation) (DA 1906/4-1 to DCD.). Work by WR, CR and RJS was supported by the Max Planck Society through funds to RJS.

## Author Contributions

RJ, MCL, CR, WR and DCD designed and performed experiments. RJ, MCL, CR and DCD analyzed the data. DCD wrote the paper with contributions from RJS and MCL.

---

**Fig. S1.**
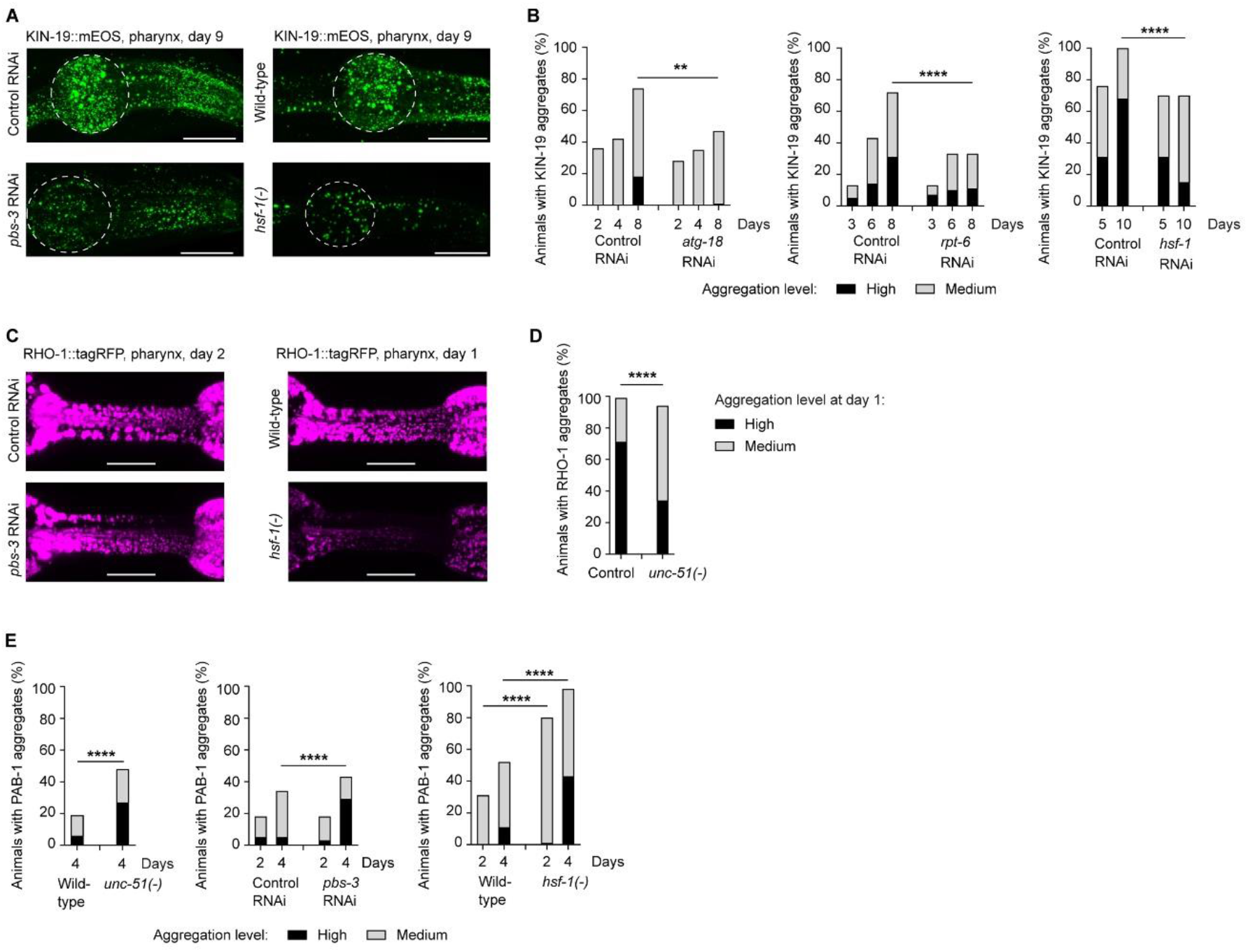
SAPA does not target stress granule aggregation-prone protein PAB-1. (A, B) Aged animals with impaired PQC have less KIN-19 aggregates in the pharynx. Representative confocal images of animals expressing KIN-19::mEOS2 in the pharynx shown as maximum z-stack projections with anterior pharyngeal bulb circled in white. Scale bar: 20 μm (A). Changes in KIN-19::mEOS2 aggregation (left panel) or in KIN-19::tagRFP aggregation (middle and right panels) evaluated over time in the population with impaired macroautophagy (*atg-18* RNAi), impaired proteasomal degradation (*rpt-6* RNAi) and reduced chaperone levels (*hsf-1* RNAi) (B). (C, D) Young animals with impaired PQC have less RHO-1 aggregates in the pharynx. Representative confocal images of RHO-1::tagRFP in the pharyngeal isthmus shown as maximum z-stack projections. Scale bar: 15 μm (C). Changes in RHO-1::tagRFP aggregation evaluated over time in the population with impaired macroautophagy (*unc-51(-)*) (D). (E) PQC inhibition accelerates PAB-1 aggregation in pharyngeal muscles. P-values determined by Fisher’s exact test and Chi-test. **p<0.01, ****p<0.0001 See also Data File S1 for number of animals evaluated and statistics.

**Fig. S2.**
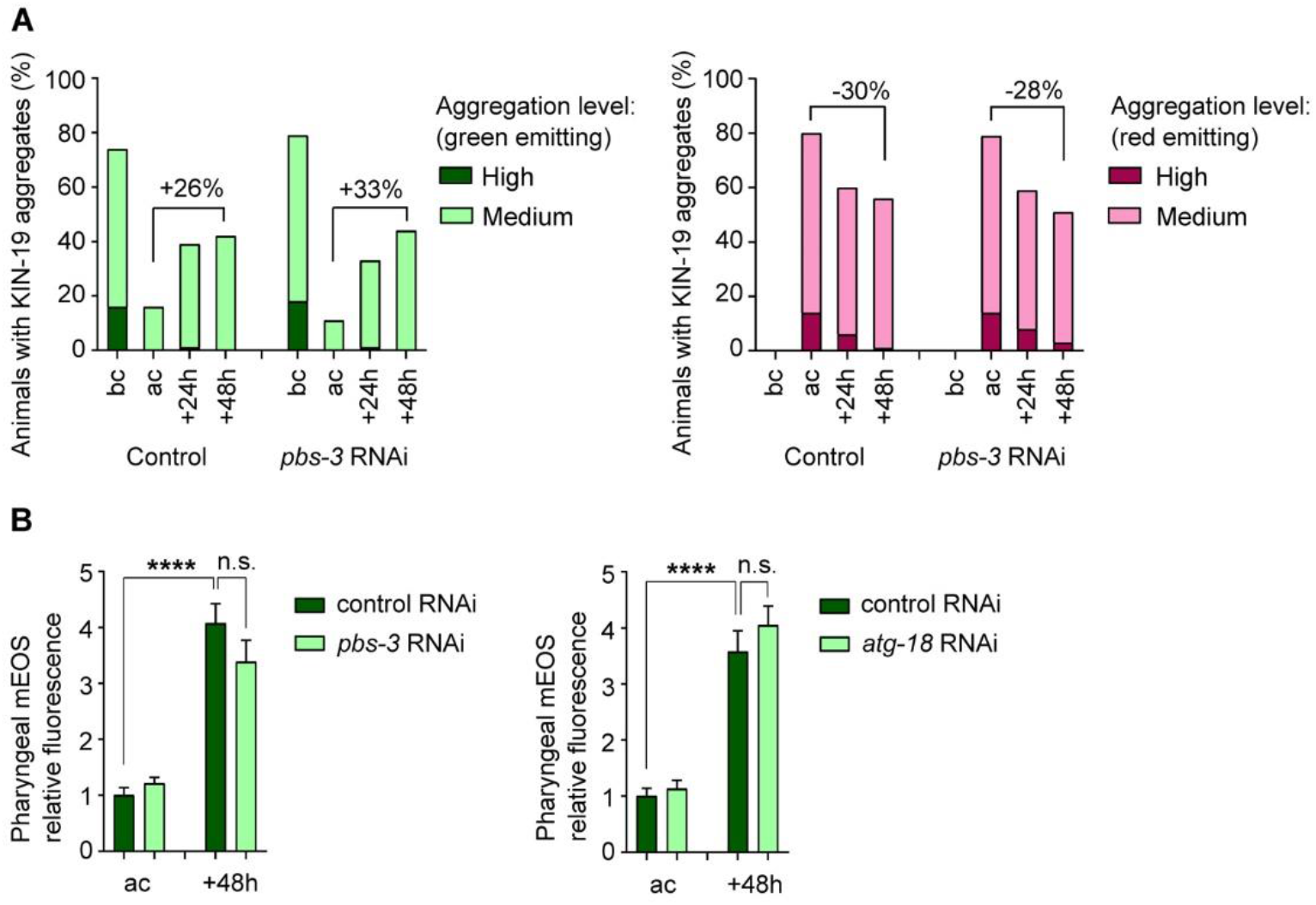
New aggregate formation in the body-wall muscles is not delayed by proteasome disruption. (A) The rate of new green-labeled KIN-19::mEOS2 aggregate formation in the body-wall muscles is similar in control and proteasome disruption conditions (*pbs-3* RNAi). Conversion was done at day 2. Before conversion (bc), after conversion (ac). In graph, difference in percent of animals with aggregation between 48h and after conversion. (B) After photoconversion of pharyngeal mEOS2 alone in young day 2 adults, monitoring of total green fluorescence shows accumulation of newly synthesized fluorescent tag in both control treatment and upon proteasome impairment (left panel, *pbs-3* RNAi) or macroautophagy impairment (right panel, *atg-18* RNAi). Relative fluorescence represents total fluorescence of treatment normalized to control RNAi after conversion. SEM shown. P-values determined by unpaired T-test (D). ****p<0.0001 See also Data File S1 for number of animals evaluated and statistics.

**Fig. S3.**
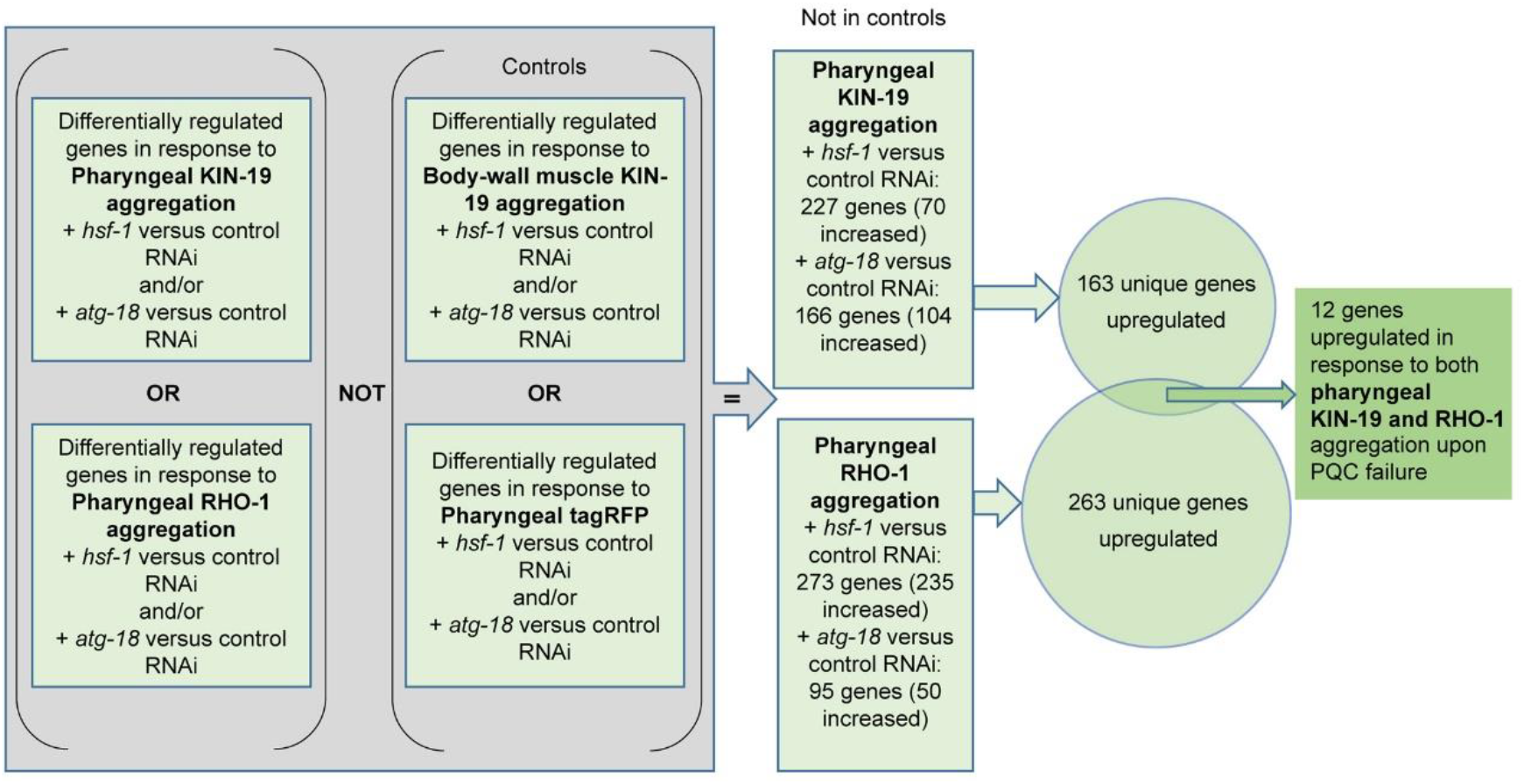
Flowchart for the analysis of RNA sequencing data. Steps to identify genes upregulated in response to PQC disruption and protein aggregation in the pharynx but not in response to PQC disruption and protein aggregation in the body-wall muscles or with the fluorescent marker alone. See also Data File S2.

**Fig. S4.**
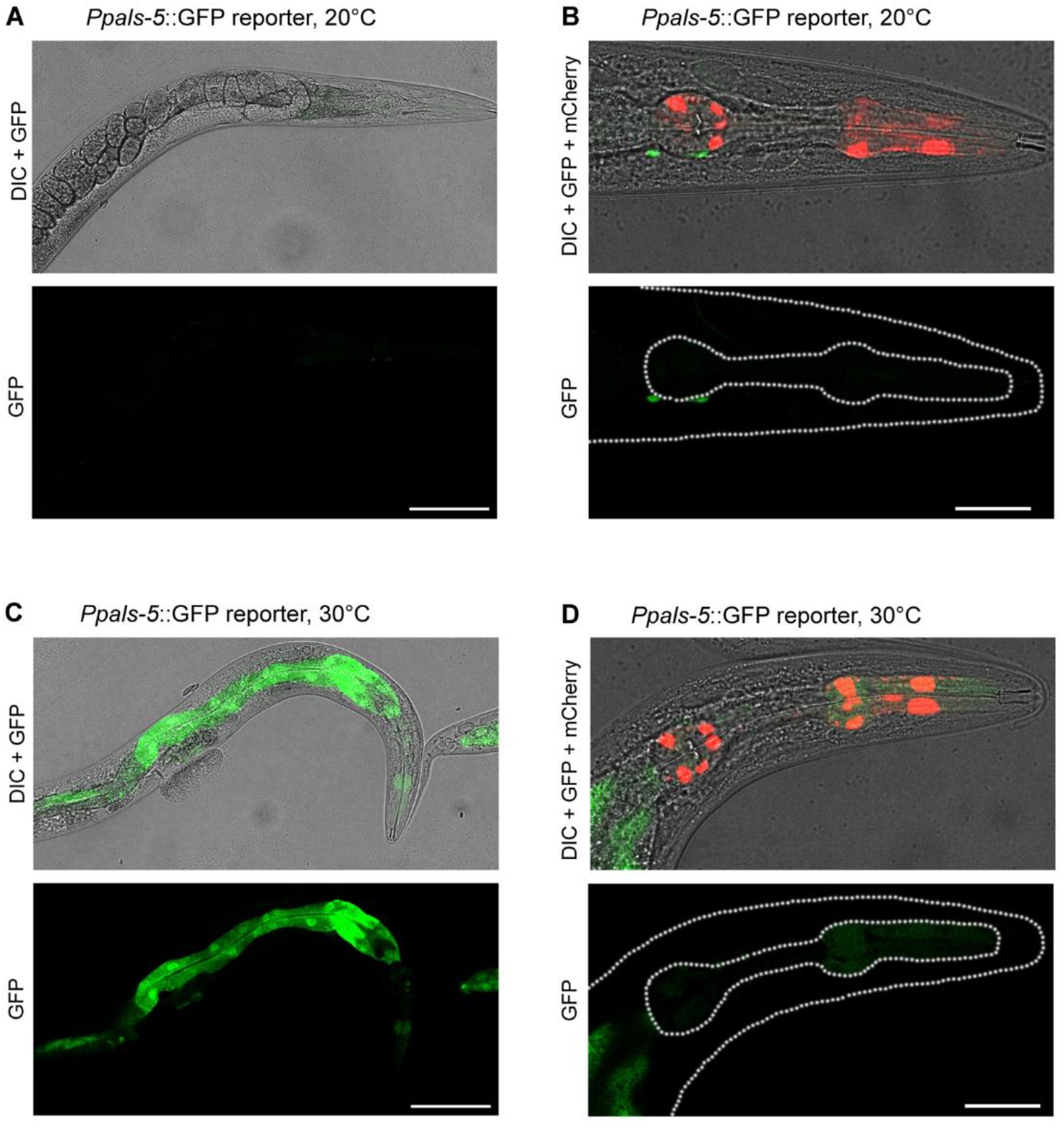
*Ppals-5::GFP* expression is induced in the pharynx and intestine. (A-D) Heat stress at 30°C for 24h induces *Ppals-5::GFP* expression in the pharynx and intestine (C, D) compared to standard growth conditions at 20°C (A, B). (D) Magnification of the head region (outlined) shows localization of GFP induced by heat stress in the pharynx (*Pmyo-2::mCherry* in top image and outlined in bottom image). Single plane representative confocal images of day 1 adults are shown. DIC: differential interference contrast. Scale bar: 100 μm (A, C), 30 μm (B, D).

**Table S1.** Genes upregulated in response to PQC disruption and protein aggregation in the pharynx

| Gene | Description | Human ortholog | Up-regulated by microsporidia* | Pharyngeal KIN-19 aggregation | Pharyngeal RHO-1 aggregation | Significantly restores RHO-1 aggregation in *hsf-1(-)* |
| --- | --- | --- | --- | --- | --- | --- |
| *pals-5* | Protein containing ALS2cr12 signature |  | yes | up with *hsf-1* RNAi | up with *hsf-1* RNAi | 3/3 repeats |
| C01B10.4 | type-B carboxylesterase | Butyryl-cholinesterase (E-value: 4e-30) |  | up with *atg-18* RNAi | up with *hsf-1* RNAi | 3/3 repeats |
| C53A5.11 | kelch domain | Actin-binding protein IPP (E-value: 1.2e-27) | yes | up with *hsf-1* RNAi | up with *hsf-1* RNAi | 3/3 repeats |
| C30H6.12 |  |  |  | up with *atg-18* RNAi | up with *hsf-1* RNAi | 3/3 repeats |
| *fbxa-30* | F-box A protein |  |  | up with *atg-18* RNAi | up with *hsf-1* RNAi | 2/3 repeats |
| F26F2.4 |  |  | yes | up with *hsf-1* RNAi | up with *hsf-1* RNAi | 2/3 repeats |
| *math-39* | MATH (meprin-associated Traf homology) domain containing |  |  | up with *hsf-1* RNAi | up with *hsf-1* RNAi | 1/3 repeats |
| T05B11.1 | F-box |  |  | up with *atg-18* RNAi | up with *hsf-1* RNAi | 1/4 repeats |
| *fbxa-86* | F-box A protein |  | yes | up with *hsf-1* RNAi | up with *hsf-1* RNAi | N.A. |
| Y105C5A.1270 |  |  |  | up with *hsf-1* RNAi | up with *hsf-1* RNAi | N.A. |
| M01G12.7 |  |  | yes | up with *hsf-1* RNAi | up with *hsf-1* RNAi | N.A. |
| K06A9.1 |  |  |  | up with *atg-18* RNAi | up with *hsf-1* RNAi | N.A. |

\*Bakowski et al. PLoS Pathog., 2014.

**Data file S1. Source data**

Excel file with source data for figures.

**Data file S2. Differentially expressed genes upon PQC failure**

Excel file, sheet 1 shows all differentially expressed genes, sheet 2 shows genes significantly regulated in transgenics with pharyngeal KIN-19 or RHO-1 aggregation and not in controls, sheet 3 shows SAPA candidates upregulated in both transgenics with pharyngeal KIN-19 and RHO-1 aggregation and sheet 4 shows enrichment analysis of SAPA components regulated by microbes.

**Data file S3. Investigation of SAPA components by RNAi**

Excel file, sheet 1 shows results of four repeats in *hsf-1(-)* background and sheet 2 shows results of two repeats in wild-type background.

